# T7 RNA polymerase-independent expression of reporter genes from a T7 promoter-driven SARS-CoV-2 replicon-encoding DNA in human cells

**DOI:** 10.1101/2024.03.01.582915

**Authors:** Ronja Friedhoff, Ghada Elfayres, Natacha Mérindol, Isabel Desgagné-Penix, Lionel Berthoux

## Abstract

Replicons, derived from RNA viruses, are genetic constructs retaining essential viral enzyme genes while lacking key structural protein genes. Upon introduction into cells, the genes carried by the replicon RNA are expressed, and the RNA self-replicates, yet viral particle production does not take place. Typically, RNA replicons are transcribed *in vitro* and are then electroporated in cells. However, it would be advantageous for the replicon to be generated in cells following DNA transfection instead of RNA. In this study, a bacterial artificial chromosome (BAC) DNA encoding a SARS-CoV-2 replicon under control of a T7 promoter was transfected into HEK293T cells engineered to functionally express the T7 RNA polymerase (T7 RNAP). Upon transfection of the BAC DNA, we observed low, but reproducible expression of reporter proteins GFP and luciferase carried by this replicon. Expression of the reporter proteins required linearization of the BAC DNA prior to transfection. Surprisingly, however, expression occurred independently of T7 RNAP. Gene expression was also insensitive to remdesivir treatment, suggesting that it did not involve self-replication of replicon RNA. Similar results were obtained in highly SARS-CoV-2 infection-permissive Calu-3 cells. Strikingly, prior expression of the SARS-CoV-2 N protein boosted expression from transfected SARS-CoV-2 RNA replicon but not from the replicon BAC DNA. In conclusion, transfection of a large DNA encoding a coronaviral replicon led to reproducible replicon gene expression through an unidentified mechanism. These findings highlight a novel pathway toward replicon gene expression from transfected replicon cDNA, offering valuable insights for the development of methods for DNA-based RNA replicon applications.

## Introduction

In late 2019, the severe acute respiratory syndrome related coronavirus 2 (SARS-CoV-2) emerged in Wuhan, China, causing the global COVID-19 pandemic. COVID-19 is associated with mild cold symptoms to severe pneumonia which can lead to hospitalization and death (1). The disease can also transform into a chronic disease known as long COVID, which can involve increased fatigue, organ injury and an increased risk for type 2 diabetes (2). As of February 2024, there had been nearly 775 million confirmed cases of SARS-CoV-2 infections worldwide with a death toll of over 7 million (source: World Health Organization). The first vaccines against SARS-CoV-2 were made available in a matter of months after the beginning of the pandemic. COVID-19 vaccine campaigns quickly proved to be especially effective in preventing severe infection (3). The search for antiviral compounds to prevent infection or complications in high-risk individuals continues to be of high importance (4). Currently, several antiviral therapies have been approved, with the SARS-CoV-2 protease inhibitor nirmatrelvir (Paxlovid) probably being the most frequently prescribed. However, the global use of antiviral compounds to target SARS-CoV-2 has been low, which is explained by a combination of disappointing effectiveness and high cost (5).

The discovery and characterization of novel putative antiviral drugs for SARS-CoV-2 has been complicated by two factors. First, SARS-CoV-2 has long been a biosafety level 3 pathogen, though an increasing number of countries are now classifying it as a level 2 pathogen (6). Secondly, SARS-CoV-2-derived viral vector systems encoding marker proteins that facilitate screening have been slow to develop, and to this day have very low infectivity (7). To circumvent those roadblocks, viral replicons have often been used instead of the wild-type coronavirus. Replicons are non-infectious subgenomic viral RNAs in which key structural genes are eliminated, making them unable to form progeny virions (8). Self-copying of the replicon RNA is still possible because all non-structural genes required for genomic replication and transcription are present. Transfection of the replicon RNA into different cell lines thus leads to expression of viral genes as well as any marker inserted in the replicon. This potentially allows the screening for antiviral compounds that target viral replication *in cellulo* (8). Prior to SARS-CoV-2, replicon systems had been created for other coronaviruses, such as SARS-CoV (sometimes called SARS-CoV-1) as well as the Middle East respiratory syndrome-related coronavirus (MERS-CoV) (9, 10).

Traditionally, replicons are introduced into mammalian cells through *in vitro* transcription of replicon cDNA into RNA, followed by transfection by electroporation. Replicon-encoded proteins are then translated by the host cell’s ribosomes, leading to RNA replication. This method, while widely employed, is plagued with drawbacks such as high cost, time-intensive procedures, and inefficiency in achieving successful electroporation of replicons into cells. Establishing cell lines supporting the stable self-replication of replicon RNAs is also not trivial. A transformative alternative is the direct expression of replicon RNA from a DNA construct within cells, offering several advantages for reverse genetics as applied to RNA viruses: (i) amplifying and purifying DNA is inexpensive compared to RNA; (ii) DNA transfection is less expensive and more efficient, routinely reaching >90% in some cellular models; (iii) stably maintaining a DNA construct is a reachable goal, through its integration into the cellular genome. Prior attempts at such an objective have been made, for instance by using a CMV promoter to govern replicon expression (11). In this study, we explored the possibility to produce SARS-CoV-2 replicon RNA from a T7 promoter-driven BAC construct introduced into human cells by standard transfection protocols. Here, we specifically report on the observed expression of two non-viral markers that are part of the replicon.

Our findings contribute to advancing DNA-based RNA replicon applications with potential implications for efficient and cost-effective reverse genetics studies in RNA virus research.

## Materials and methods

### Plasmids

The SARS-CoV-2 N protein-expressing plasmid pEZY3-N was described before (12). T7-CMVtrans-FFLuc-polyA (13), which expresses the firefly luciferase under control of the T7 promoter, was a gift from Marcel Bruchez (Addgene plasmid #101156). The codon-optimized T7RNAP coding sequence was amplified from T7 opt in pCAGGS (14) (Addgene #65974; a kind gift from Benhur Lee) by PCR. PCR reactions were done in 50 µl total volume, using 0.5 µl of Q5 HF DNA polymerase (New England Biolabs), 10 µM of dNTPs (New England Biolabs) and 10 µM of primer XhoI-T7-5’ [AGCTCTCGAGACCATGAACACCATCAATATTGCC] and EcoRI-§-T7-3’ [GCATGAATTCTCAGGCAAATGCGAAATCGGA]. Amplification conditions were as follows: 20 sec denaturation at 98°C, 25 cycles (10 sec at 98°C, 20 sec at 58°C, 2 min at 72°C) and 5 min at 72°C to complete synthesis. PCR products were purified using the QIAquick purification kit (QIAGEN), then cut with XhoI and EcoRI, and finally gel-purified using the QIAEX II kit (QIAGEN). The purified inserts were ligated in separate reactions into pLPCX(AB) and pMIP (15) that were linearized by digestion with XhoI and EcoRI. pLPCX(AB) is a version of pLPCX (Clontech Laboratories) in which the EcoRI-ClaI region of the multi-cloning site was removed and replaced by a duplex created by annealing oligodeoxynucleotides Linker-EcoBamNotCla-s (5’-AATTCACGGATCCTTGCGGCCGCAT) and Linker-EcoBamNotCla-as (5’-CGATGCGGCCGCAAGGATCCGTG). Ligation reactions were performed in 20 µL total volume, with 2 µg vector, half of a purified reaction product, and 1 µL of T4 DNA ligase (New England Biolabs). Following ligation, 1:10 vol. of the ligated product was transformed into DH5α *E. coli* by electroporation. Clones were analyzed by restriction enzyme digestions followed by Sanger sequencing.

### Cell culture

Human embryonic kidney (HEK)293T cells were maintained in DMEM (Hyclone) supplemented with 10% fetal bovine serum (Hyclone) and penicillin-streptomycin (Hyclone). HEK293T cells lentivirally transduced with SARS-CoV-2 N have been described before (12). The lung adenocarcinoma epithelial cells Calu-3 were maintained in EMEM (Hyclone) supplemented with 10% fetal bovine serum.

### Retroviral vector production and HEK293T transductions

To create HEK293T cells stably expressing T7 RNAP, two retroviral vectors were used. HEK293T cells plated at approximately 80% confluence in 10-cm plates were polyethylenimine (PEI) transfected with pLPCX(AB)-T7RNAP (10 µg) along with psPAX2 (10 µg) and pMDG (5 µg) as described before (16). To generate the control vector, cells were transfected with the empty vector pLPCX(AB) instead of pLPCX(AB)-T7RNAP. We also generated MIP-based vectors (15) by transfecting HEK293T cells with pMIP-T7RNAP (10 µg), pCL-Eco (10 µg) and pMDG (5 µg) as described before (17). Again, a control vector was produced by transfecting with the empty pMIP instead of pMIP-T7RNAP. The following day, supernatants were removed and replaced with complete medium. Two days post transfection, supernatants were harvested, clarified by low-speed centrifugation and aliquoted. HEK293T cells plated at 70% confluency in 10-cm plates were transduced by adding 5 mL of vector-containing supernatant. The next day, supernatants were replaced with complete medium. Two days post transduction, all transduced cells were placed in 1 µM puromycin (Gibco). Selection was allowed to proceed for 4 to 9 days; control untransduced cells were killed by the treatment.

### Luciferase-expressing plasmid DNA transfection and luciferase assays

Cells plated in 6-well plates at 70 % confluency were transfected with 2 µg per well of T7-CMVtrans-FFLuc-polyA using PEI. Cell supernatants were replaced with fresh medium six hours later. Cells were processed for luciferase assay approximately 36 h later. The Steady-Glo Luciferase Assay System kit from Promega was used for all luciferase assays. Cells grown in 6-well plates were washed with PBS, trypsinized and pelleted by centrifugation. Post-centrifugation pellets were resuspended in 300 µL of complete medium and plated in three wells of a 96-well plate (black wall, clear bottom) at 100 µL per well. 100 µL of Steady-Glo Reagent were immediately added to each well. Luciferase activity was measured using the Biotek Synergy HT microplate reader according to the manufacturer’s instructions.

### Preparation of SARS-CoV-2 replicon-encoding bacterial artificial chromosome (BAC) DNA and *in vitro* transcription of replicon RNA

The pSMART BAC v2.0 vector containing the SARS-CoV-2 (Wuhan-Hu-1) non-infectious replicon (18) was obtained from BEI resources (#NR-54972). To transform Stbl3 *E coli* cells (Invitrogen) with the BAC construct, one vial of One Shot Stbl3 chemically competent cells was thawed on ice and mixed with 5 µL of DNA. The mixture was incubated on ice for 30 min followed by heat shock treatment at 42 °C for 45 sec without shaking. The vial was placed back on ice for 2 min, then 250 µL of pre-warmed SOC medium was added followed by incubation at 37°C for 1 h. The cells were plated onto LB plates containing 20 μg/mL chloramphenicol, and then cultured overnight at 37 °C. Individual colonies were isolated and grown, and then the BAC DNA was purified using the PhasePrep BAC DNA Kit (Sigma Aldrich) according to the manufacturer’s instructions, and analyzed by restriction enzyme digestion.

For *in vitro* transcription, the BAC replicon-encoding DNA was first linearized with SwaI (New England Biolabs), and then purified by phenol/chloroform extraction and ethanol precipitation. The fragments were dissolved in nuclease-free water. The mMESSAGE mMACHINE T7 transcription kit (Invitrogen) was used to generate the replicon RNAs in the correct orientation from the linearized vector according to manufacturer’s instruction. Briefly, 180 μL of T7 transcription reaction, containing 7.2 μg of linearized BAC, T7 RNAP as well as GTP, was incubated at 37 °C for 2.5 h. After incubation, 9 μL of TURBO DNase was added and the reaction was incubated at 37 °C for 15 min to digest DNA. The resulting RNA was purified using the Monarch RNA cleanup kit (New England Biolabs) and analyzed by agarose gel electrophoresis.

### SARS-CoV-2 replicon BAC DNA transfections

Cells plated in 6-well plates were transfected at 50 % confluency with various amounts of BAC DNA, either intact or linearized with SwaI immediately prior to transfections. Following SwaI digestions (2 h at 25 °C), BAC DNA was heated for 20 min at 65 °C to denaturate the enzyme. Transfections were done using either Lipofectamine 3000 (Invitrogen), PEI or electroporation. For Lipofectamine 3000 transfections, 2 µg of BAC DNA in 125 µl Opti-MEM (Gibco) containing 3 µl of P3000 reagent were mixed with 125 µl of Opti-MEM containing 3 µl of Lipofectamine 3000.

After incubation at room temperature for 15 min, the lipofectamine:DNA mixture was added onto the cells. PEI transfections were performed as detailed previously (19), whereas electroporations were performed exactly as for the replicon RNA (see below). In some experiments, cells were transfected with 5 μg of pEZY3-N concomitant with BAC DNA transfections. In some instances, remdesivir (Cayman Chemical) was added at a concentration of 100 nM or 1 μM, immediately after transfection and treatment was repeated with the media change the next day. Control cells were mock-transfected using identical conditions.

### SARS-CoV-2 replicon RNA transfections

HEK293T cells were harvested using Trypsin/EDTA (Thermo Fisher Scientific), washed with PBS, and resuspended in Neon Resuspension Buffer R (Invitrogen) to a final density of 1 × 10^7^ cells/mL. 10 μg of replicon RNA were added onto 1 × 10^6^ resuspended cells. The mixture was immediately transferred to Neon tips (100 µL, Invitrogen), and electroporation was carried out in the Neon Device (Invitrogen) using the cell type-specific preloaded parameters. Transfected cells were transferred immediately into 6-well plates containing prewarmed DMEM medium supplemented with 10% FBS and without antibiotics, and placed in antibiotics-containing medium the next day. In some cases, cells were electroporated with 5 μg of pEZY3-N simultaneously to replicon RNA transfections.

### Analysis of reporter proteins expression

To analyze GFP expression by flow cytometry, cells in a 6-well plate were trypsinized with 500 µl trypsin and the reaction was stopped with 500 µl of complete media. 800 µl of each cell suspension were then fixed using 4% formaldehyde diluted in PBS. The percentage of GFP-positive cells was determined by analyzing 20,000 cells on a FC500 MPL cytometer (Beckman Coulter) using the CXP Software (Beckman Coulter), or CytoFlex S (Beckman Coulter). Flow cytometry data were analyzed using FlowJo (Becton Dickinson) or FCS Express (De Novo Software). To analyze luciferase expression, 2 x 100 µL of the cell suspension were processed for firefly luciferase quantification assay, as described above.

### Fluorescence microscopy

GFP expression in live cells was examined using an Axio Observer inverted fluorescence microscope (Zeiss) with a 20X objective, two days post transfection with SARS-CoV-2 replicon BAC DNA or with replicon RNA. Images were recorded using the ZEN software.

## Results

### Transduction of functional T7 RNAP into HEK293T cells

We generated two retroviral vector constructs, LPCX(AB)-T7RNAP and MIP-T7RNAP, and used them to transduce HEK293T cells followed by elimination of non-transduced cells by puromycin treatment. “Empty” LPCX(AB) and MIP vectors were used as controls. Validating T7 RNAP expression by Western blotting was not possible due to the unavailability of antibodies. However, we were able to perform a functional assay for T7 RNAP, by transfecting the transduced cell populations generated with a construct expressing the firefly luciferase under control of the T7 promoter (Fig 1). We observed strongly increased luciferase activity in cells transduced with LPCX(AB)-T7RNAP or MIP-T7RNAP compared to the background levels of luminescence activity in cells transduced with the empty vector controls (Fig 1). These results demonstrate the functionality of T7 RNAP in the HEK293T cells generated.

**Fig 1.**
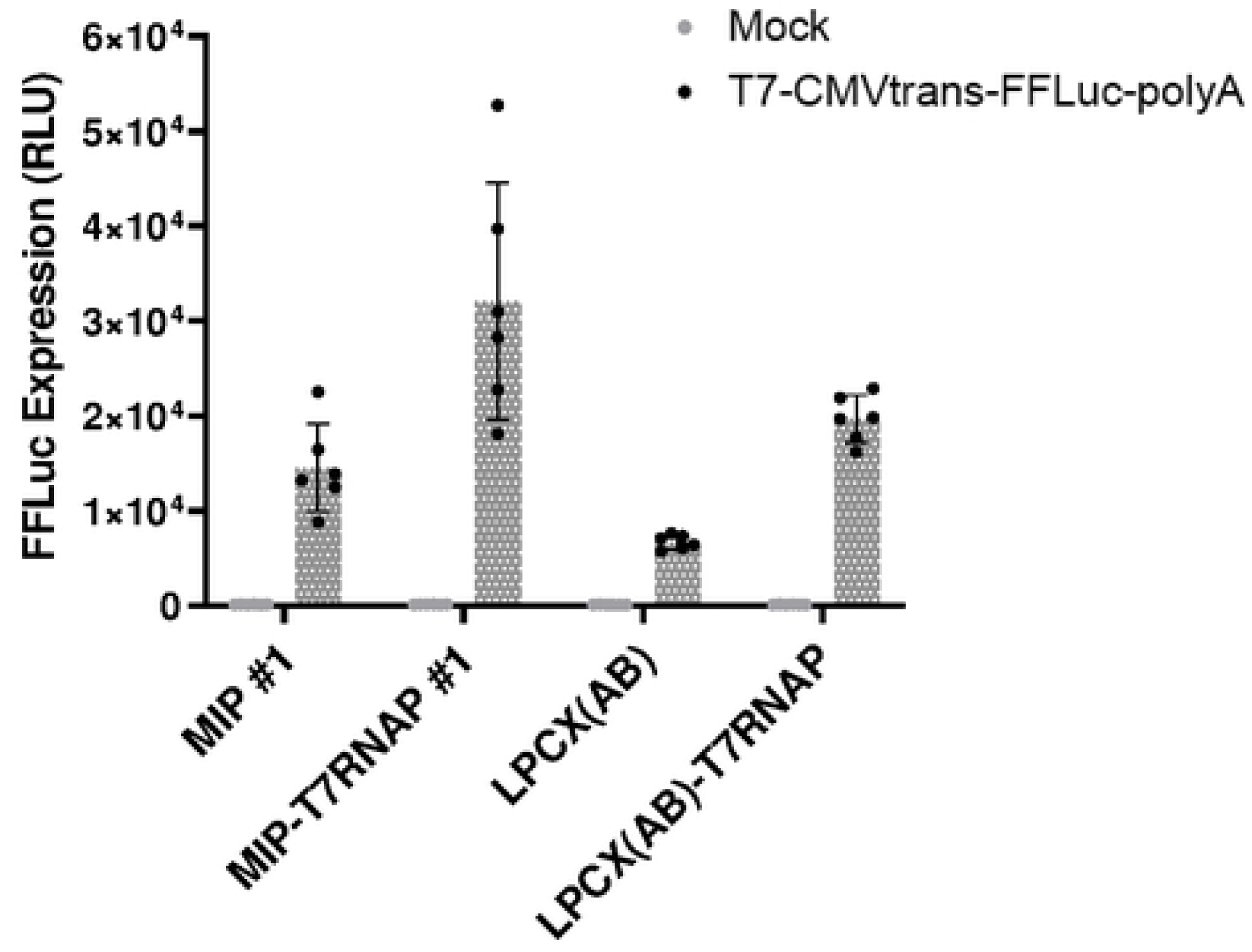
HEK293T cells transduced with T7 RNAP support T7 promoter-dependent expression. HEK293T cells were transduced with MIP-T7RNAP or LPCX(AB)-T7RNAP or with the empty vectors as controls. Cells were treated with antibiotics to kill untransduced cells and then were transfected in triplicates with a plasmid expressing luciferase under control of the T7 promoter. Luciferase activity was quantified the next day as described in the Methods section. RLU, relative lights units.

### DNA linearization-dependent, T7 RNAP-independent, remdesivir-insensitive expression of replicon-encoded GFP following BAC DNA transfection in HEK293T cells

The SARS-CoV-2-encoding BAC construct used in this project has deletions in the viral structural proteins S, E and M, and the coding sequences for the firefly luciferase and GFP are expressed as a fusion protein in place of S (18). In order to explore conditions permitting the *in cellulo* transcription and replication of the SARS-CoV-2 replicon, we used a PEI-based protocol to introduce the BAC DNA into the HEK293T-LPCX(AB) cells (Fig 2A) and HEK293T-LPCX(AB)-T7RNAP (Fig 2B). Furthermore, we transfected both linearized and non-linearized SARS-CoV-2 replicon BAC DNA, with the expectation that linearization by SwaI digestion would be necessary for the transcription of replicon RNA. Finally, transfection of the linearized BAC DNA in the HEK293T-LPCX(AB)-T7RNAP was also done in presence of remdesivir, a nucleoside analog that abrogates replicon RNA replication (20). GFP expression was measured 2 days later by flow cytometry. The data obtained show expression of replicon-encoded GFP in both empty vector-transduced cells (Fig 2A) and T7RNAP-expressing cells (Fig 2B) upon transfection with linearized BAC DNA. The observed percentages of GFP expression were low but non-negligible (in the 0.4-0.8% range) in both cell populations. In contrast, little to no GFP signal was detected in cells transfected with non-linearized SARS-CoV-2 replicon, showing that linearization was essential for GFP expression from this DNA. Finally, treatment with 100 nM remdesivir did not prevent GFP expression (Fig 2B, bottom left panel).

**Fig 2.**
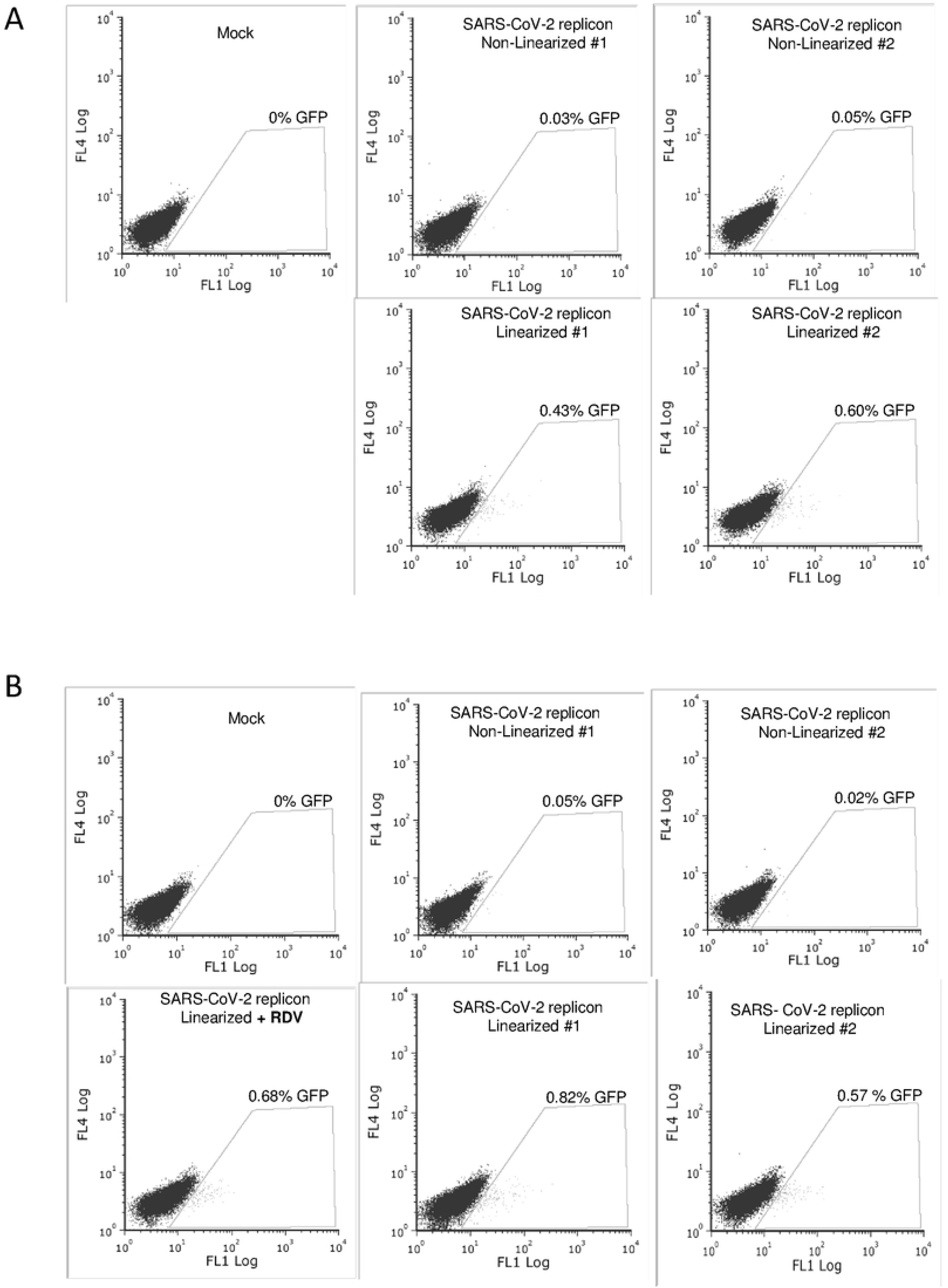
PEI transfection of SARS-CoV-2 replicon DNA leads to GFP expression that is dependent on DNA linearization but not T7 RNAP and is remdesivir-insensitive. HEK293T cells transduced with the empty retroviral vector LPCX(AB) (A) or with LPCX(AB)-T7RNAP (B) were PEI-transfected with the pSMART-BAC-T7-scv2 replicon or were mock-transfected as a control. BAC DNA was linearized or not with SwaI immediately prior to transfections. Two independent replicon DNA transfections were done for both non-linearized and linearized replicon DNA. For the linearized replicon, an additional transfection was performed in the presence of 100 nM remdesivir (bottom left dot plot). GFP expression was analyzed two days post-transfection by flow cytometry.

The BAC DNA utilized in this study, with a substantial length of approximately 36 Kbp, poses a challenge for transfection, given its size. To address this, we repeated the experiment using Lipofectamine 3000 to introduce linearized or non-linearized replicon BAC DNA into HEK293T-LPCX(AB) (Fig 3A) and HEK293T-LPCX(AB)-T7RNAP (Fig 3B). Remarkably, GFP expression was consistently observed in both cell populations following transfection of linearized BAC DNA, but not non-linearized DNA. GFP was detected in 0.4-0.5% cells in both cell populations, and once again, remdesivir treatment had no effect. We used fluorescence microscopy to record images of transfected HEK293T-LPCX(AB)-T7RNAP cells in the absence (S1 Fig) or the presence (S2 Fig) of remdesivir. Fluorescence was relatively weak, as expected from the flow cytometry results, but fluorescent cells were adherent and appeared not to be at an advanced apoptotic state that may induce unspecific autofluorescence. The absence of autofluorescence artifacts is also evidenced by the fact that fluorescence was recorded in the FL1 channel of the cytometer, but not other channels like FL4 (Fig 2, Fig 3), proving that fluorescence was associated with GFP expression. Altogether, the data presented in Fig 2 and Fig 3 show that GFP expression following replicon BAC DNA is linearization-dependent, but T7 RNAP-independent and remdesivir-insensitive. These observations point to GFP expression in the absence of full-length replicon RNA expression or replication.

**Fig 3.**
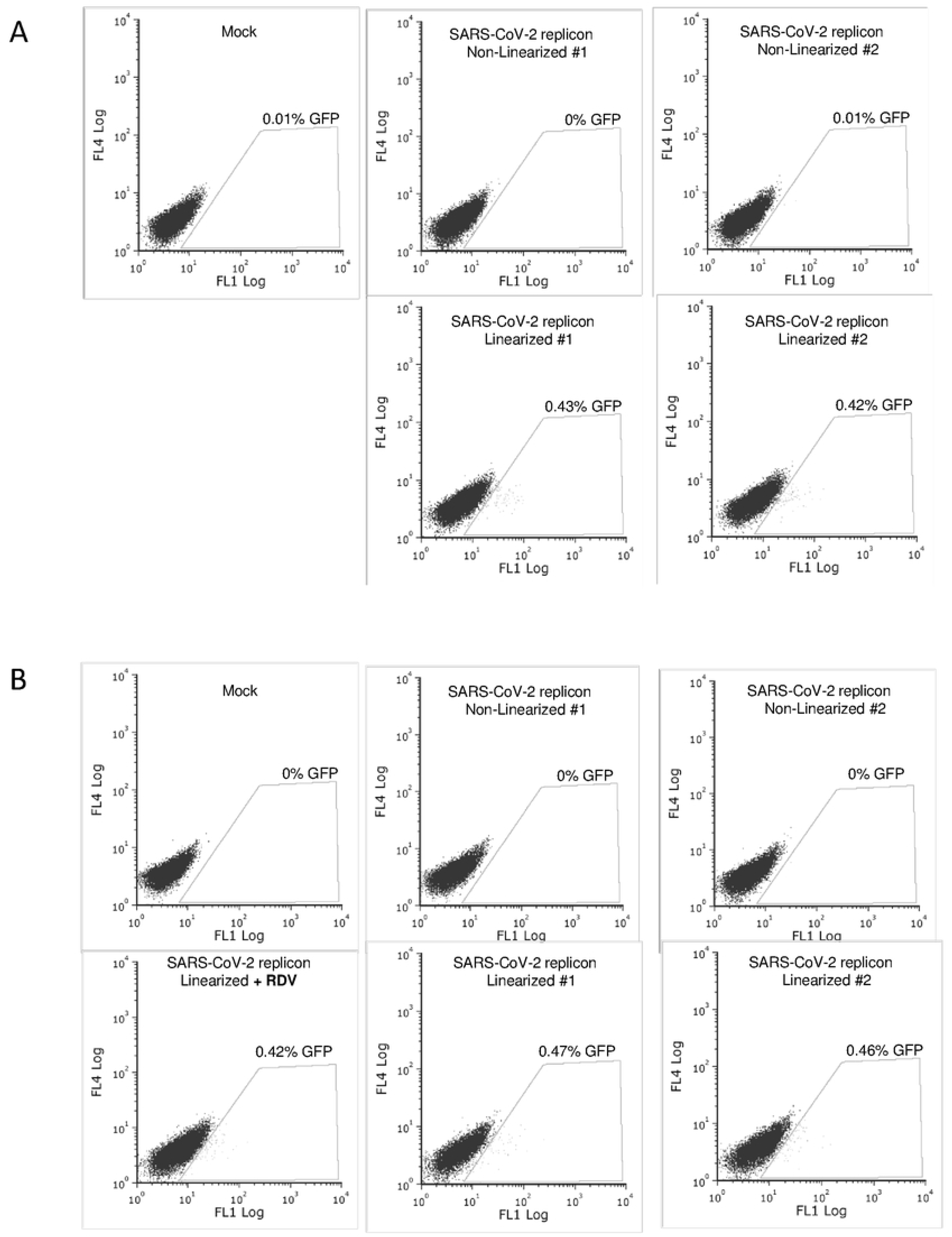
Lipofectamine transfection of SARS-CoV-2 replicon DNA leads to GFP expression that is dependent on DNA linearization but not T7 RNAP and is remdesivir-insensitive. HEK293T transduced with LPCX(AB) (A) or with LPCX(AB)-T7RNAP (B) were then transfected with pSMART-BAC-T7-scv2 DNA, linearized or not, and in the absence or presenct of remdesivir, exactly as in Fig. 2, except that transfections were performed using Lipofectamine 3000. Cells were analyzed by flow cytometry two days later.

### Luciferase activity following SARS-CoV-2 replicon-encoding DNA transfection

The replicon used in this study encodes a firefly luciferase-GFP fusion protein. Thus, GFP expression is expected to correlate with luciferase activity. HEK293T cells transduced with LPCX(AB) or LPCX(AB)-T7RNAP were transfected with SwaI-linearized or non-linearized replicon BAC DNA, and luciferase activity was measured in cellular lysates (Fig 4). The data obtained show an absence of luciferase activity upon transfection of non-linearized DNA, since RLU values were similar to background levels observed in mock-transfected cells. Luciferase activity was approximately 1.8-fold over background levels upon transfection of linearized DNA, and luciferase activity levels were similar in both LPCX(AB) and LPCX(AB)-T7RNAP cells. Furthermore, remdesivir treatment did not affect luciferase activity levels (Fig 4). Thus, these results recapitulate the GFP expression data: linearization of the SARS-CoV-2 replicon DNA is required for luciferase expression, but T7 RNAP is not, and expression is remdesivir-insensitive.

**Fig 4.**
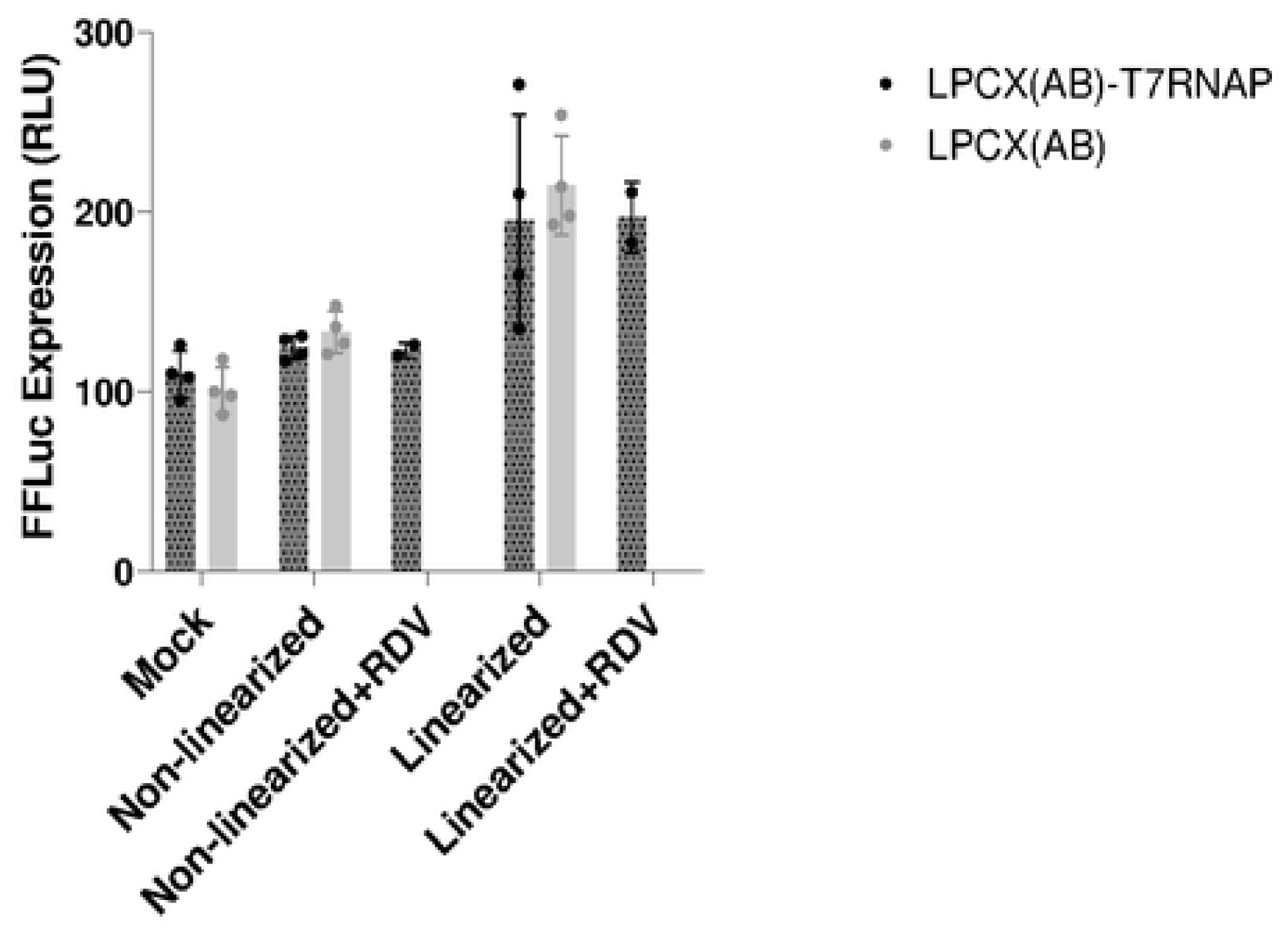
SARS-CoV-2 replicon DNA-encoded luciferase expression is dependent upon DNA linearization but occurs in the absence of T7 RNAP and is insensitive to remdesivir. HEK293T cells transduced with LPCX(AB) or with LPCX(AB)-T7RNAP were PEI-transfected with pSMART-BAC-T7-scv2, linearized or not as indicated, and treated or not with remdesivir (RDV). Luciferase activity was quantified two days later as detailed in the Methods section, and is expressed in relative light units (RLU). Shown are averaged values from duplicate transfections with two luciferase assays done for each transfection.

### Characterization of SARS-CoV-2 replicon in HEK293T cells following replicon DNA or RNA transfection in presence of SARS-CoV-2 N

Prior presence of the SARS-CoV-2 N structural protein has been shown to improve replicon RNA survival and expression (11). To uncover a possible role for SARS-CoV-2 N in stimulating the T7 RNAP-independent expression of marker genes from the replicon BAC DNA, we used previously generated HEK293T cells lentivirally transduced to express the N protein (12), and we also transiently transfected a mammalian expression plasmid encoding the same protein. As shown in Fig 5A, expression of N in both transiently transfected and stably transduced cells was confirmed by Western blot. We then investigated whether GFP expression from the SARS-CoV-2 replicon RNA would be stimulated by N expression in transiently transfected cells (Fig 5B) or in lentivirally transduced cells (Fig 5C). SARS-CoV-2 RNA replicon electroporation was low (less than 1%), as expected, but GFP intensity was significantly higher than what was seen upon replicon DNA transfection. Fluorescence microscopy observations confirmed that GFP-expressing cells following RNA electroporation appeared intact and that GFP intensity was relatively high compared with BAC DNA transfections (S3 Fig). Transient transfection of N did not significantly enhance the percentage of GFP-expressing cells (Fig 5B), but this may be due to the concomitant co-transfection, instead of N being transfected prior to RNA electroporation, which was not feasible due to transfection-associated cytotoxicity. In contrast, stable N transduction greatly increased the efficiency of electroporated replicon RNA expression, as 8.1% cells were GFP-positive, a nearly 20-fold increase (Fig 5C). This was also reflected in fluorescence microscopy observations (S3 Fig). Moreover, remdesivir treatment strongly reduced replicon RNA expression (a 33-fold decrease) in these N-transduced cells (Fig 5C), indicating that replicon RNA self-replication was necessary for efficient expression.

**Fig 5.**
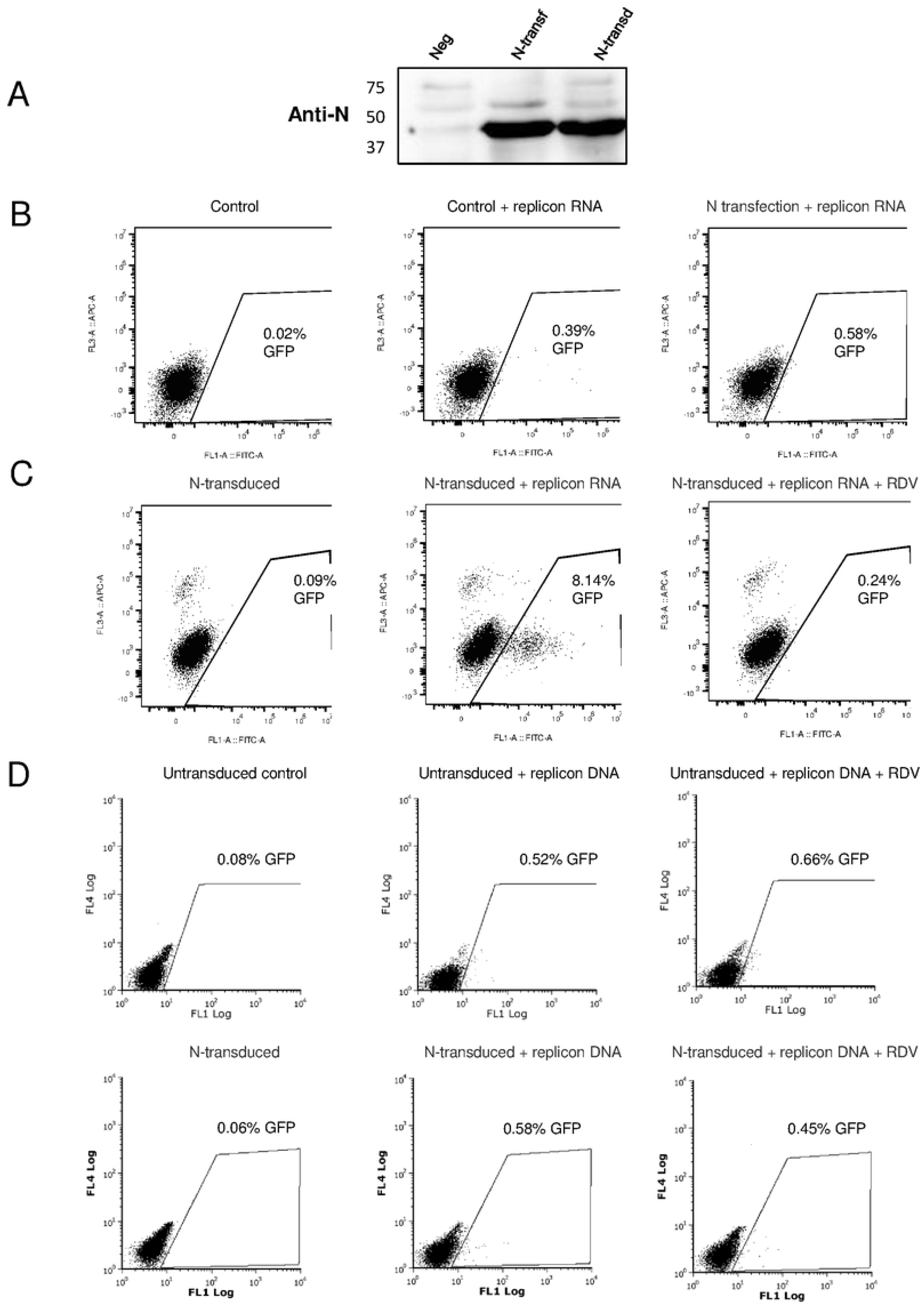
SARS-CoV-2 N promotes GFP expression following replicon RNA but not BAC DNA transfection. (A) Western blot showing N expression in HEK293T cells two days following PEI transfection of pEZY3-N, and in HEK293T cells lentivirally transduced with N. “Neg” are non-transfected, non-transduced HEK293T cells. (B) Effect of SARS-Cov-2 N transfection on SARS-CoV-2 replicon RNA expression. HEK293T cells were untransfected (Control, left dot plot), or were electroporated with SARS-CoV-2 replicon RNA (center plot), or were co-electroporated with the replicon and with SARS-CoV-2 N (right plot). Cells were analyzed for GFP expression by flow cytometry two days later. (C) Effect of SARS-CoV-2 N transduction on SARS-CoV-2 replicon RNA expression. Cells stably transduced with SARS-CoV-2 N were left untransfected (left plot), or were electroporated with SARS-CoV-2 replicon RNA in the absence (center plot) or the presence (right plot) of 1 μM remdesivir. GFP expression was analyzed by flow cytometry two days later. (D) Effect of SARS-CoV-2 N transduction on GFP expression from BAC replicon DNA. Control untransduced cells (top three plots) and N-transduced cells (bottom three plots) were left untransfected (left plots), or were electroporated with SwaI-linearized pSMART-BAC-T7-scv2 in the absence (center plots) or the presence (right plots) of 1 μM remdesivir. GFP expression was analyzed by flow cytometry two days later.

Next, we tested what effect would N transduction have on T7 RNAP-independent GFP expression following replicon DNA transfection (Fig 5D). As in previous experiments, we observed a small but significant population (around 0.5%) of GFP-positive cells following the PEI transfection of linearized replicon BAC DNA into control HEK293T cells. The proportion of GFP-expressing cells was not significantly modulated by the presence of the transduced N, or upon treatment with remdesivir.

Collectively, the results shown in Fig 5 demonstrate that T7 RNAP-independent expression of GFP from the SARS-CoV-2 replicon BAC DNA is insensitive to the presence of the viral protein N, unlike gene expression from replicon RNA. In addition, remdesivir inhibits GFP expression from transfected replicon RNA but not replicon DNA, showing that the former, but not the latter, requires replicon RNA replication.

### GFP expression from SARS-CoV-2 replicon-encoding DNA in Calu-3 cells

In order to test whether a reporter gene carried by a DNA encoding a SARS-CoV-2 replicon could be detected in a cell line relevant to SARS-CoV-2 replication, we used the pulmonary cell line Calu-3. Linearized replicon BAC DNA was electroporated (as PEI was found to be inefficient in this cell line) alone or along with a plasmid expressing T7 RNAP. As shown Fig 6, less than 1% cells were GFP-positive, similar to HEK293T cells, and the frequency of GFP-expressing cells was not improved in the presence of T7 RNAP. As seen by fluorescence microscopy, the GFP-expressing cells appeared intact, suggesting that the fluorescence was not nonspecific autofluorescence (S4 Fig). Like before, treatment with remdesivir did not inhibit GFP expression from transfected replicon BAC DNA. Thus, the results obtained in Calu-3 cells corroborate the conclusions reached in HEK293T cells.

**Fig 6.**
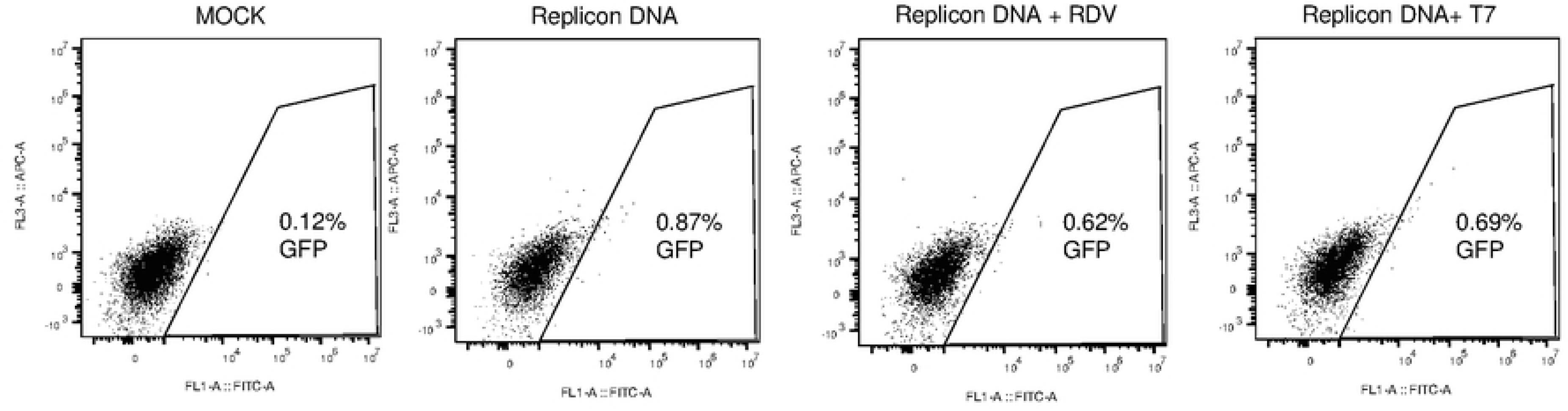
Transfection of replicon BAC DNA in Calu-3 cells yields T7 RNAP-independent and remdesivir-insensitive GFP expression. Calu-3 cells were electroporated with SwaI-linearized pSMART-BAC-T7-scv2 and co-transfected or not with T7 RNAP or in the presence of 1 μM remdesivir (RDV) as indicated. Cells were analyzed by flow cytometry two days later.

## Discussion

Biological studies of human viruses and the search for antiviral compounds require convenient methods to manipulate and mutate viral genomes and then to introduce them into human cells. For coronaviruses as well as many other RNA viruses, this can be achieved through reverse genetics (21), whereby a cDNA copy of the RNA genome is introduced into a DNA vector such as the BAC used in this study, and put under control of a microbial promoter such as the T7 RNAP-dependent promoter also used in the present study. The constructed DNA is amplified, typically in bacteria, yeast or insect cells, then purified and *in vitro* transcribed to yield a viral genomic RNA that is then introduced into mammalian cells. This viral RNA can self-replicate, thus acting as a replicon, and it can also lead to the production of novel, infectious viral particles. SARS-CoV-2 has presented additional challenges as it long was a biosafety level 3 pathogen, greatly decreasing the number of laboratories in which the wild-type virus could be studied. Thus, several groups have created reverse genetics systems to express subgenomic rather than full-length SARS-CoV-2 RNAs, as these subgenomic replicons may be used in level 2 confinement laboratories. In four different SARS-CoV-2 replicons created in 2021, the deleted viral genes included S (22), N (23), E and ORF3 (24), or the replicon used here, which is truncated for S, M and E (18). In all cases, the deleted open reading frames were replaced with marker genes such as GFP, luciferase or mCherry, allowing for the convenient measurement of replicon RNA replication and expression.

Most replicon systems rely on the *in vitro* generation of the replicon RNA followed by its introduction into cells by transfection (electroporation being the most common method used). This creates a bottleneck to upscaling, as both *in vitro* transcription and electroporation are expensive and RNA is notoriously unstable and more difficult to transfect into cells, compared with DNA. To bypass these limitations, we explored here the possibility of introducing replicon DNA, rather than RNA, into human HEK293T cells. In this scenario, expression of the replicon would occur through transcription of the replicon cDNA *in cellulo* rather than *in vitro*. Toward this aim, we constructed cells stably expressing T7 RNAP, and verified that the polymerase was indeed functional. Several teams have shown that T7-specific transcription could take place in eucaryotic cells (25), using for instance a T7 RNAP expression construct stably integrated into a mammalian cell’s genome (26). The T7 coding sequence that was used here was codon-optimized precisely to allow for higher expression levels in mammalian cells (14). However, transfection of the T7-dependent replicon-encoding BAC did not lead to T7-specific replicon expression, since we observed low levels of replicon-encoded GFP-positive cells both in the presence and absence of T7 RNAP. GFP detection in ⁓0.5% of the cells was reproducible and independent of the method used for transfecting cells. However, it was not due to background fluorescence nor was it an artifact of cytotoxicity, since fluorescence was not observed when the BAC DNA was not linearized, and fluorescent cells did not show obvious signs of cytotoxicity when observed by microscopy. Moreover, luciferase expression was similarly detected in a T7 RNAP-independent, linearization-dependent fashion. We conclude that introduction of the replicon-encoded BAC DNA into HEK293T cells can lead to expression of replicon genes through an unidentified mechanism, an unexpected finding that warrants further investigation.

What mammalian polymerase could direct the expression of GFP/luciferase from the transfected BAC DNA, in the absence of a mammalian promoter on the BAC construct? One clue might emerge from the fact that linearization was required. BAC DNA is supercoiled (27), similar to common plasmids. Thus, perhaps the unknown polymerase mobilized here does not accommodate supercoiled DNA of bacterial origin. However, a simpler explanation would be that gene expression requires RNA transcription termination which is allowed by the linearization step, similar to T7 RNAP-led transcription. We did not attempt to analyze the RNA or RNAs expressed from the BAC DNA, and thus it is not clear whether the mRNA for GFP and luciferase is identical to the SARS-CoV-2 genomic RNA synthesized by T7 RNAP (full-length), or whether shorter mRNAs are synthesized. The latter would indicate that a cryptic promoter is present in the SARS-CoV-2 genome, whereas the former would suggest that *cis*-acting elements present to the 5’ of the replicon sequence are involved in the transcription. Interestingly, GFP expression was insensitive to remdesivir, and was not improved by the concomitant presence of SARS-CoV-2 N, a protein known to promote replication of the replicon RNA (28, 29), as also observed in this study. These two observations suggest that the mRNA encoding GFP and luciferase is not self-replicating. In turn, this suggests that it is in fact a subgenomic RNA produced from a cryptic promoter present in the SARS-CoV-2 genome.

At the moment, it is not clear whether the BAC DNA T7 promoter is functional or not in HEK293T cells expressing T7 RNAP, as the absence of detectable replicon RNA might be due to degradation rather than a lack of synthesis. We are currently developing further a DNA-based SARS-CoV-2 replicon system, that will complement systems that have been described recently (11).

## Conclusions

*In vitro* RNA transcription-free reverse genetics systems are desirable for coronaviruses and other RNA viruses. In the process of developing protocols for *in cellulo* replicon RNA expression from a large cDNA plasmid, we came across the unexpected observation that reporter genes were expressed in the apparent absence of replicon RNA replication. Despite low levels of reporter protein expression, it was found to be highly reproducible, required linearization of the plasmid, and occurred through an undetermined transcription mechanism. This observation is relevant to the development of novel tools for RNA virus research, including replicons and viral vectors.

## Acknowledgements

We are grateful to Marcel Bruchez (Carnegie Mellon University, Pittsburgh, PA) and Benhur Lee (Icahn School of Medicine, New York City, NY) for sharing plasmid DNAs through Addgene, as well as Dai Wang and Merck & Co (West Point, PA) for sharing the SARS-CoV-2 replicon-encoding BAC construct through BEI Resources, NIAID, NIH. We also thank Andres Finzi (Université de Montréal) for providing the Calu-3 cells.

## Supporting information

**S1 Fig**. Fluorescence microscopy analysis of BAC DNA-transfected HEK293T cells

**S2 Fig**. GFP expression in BAC DNA-transfected HEK293T cells in presence of remdesivir

**S3 Fig**. Fluorescence microscopy analysis of replicon RNA-transfected HEK293T cells transduced or not with SARS-CoV-2 N

**S4 Fig**. GFP expression in BAC DNA-transfected Calu-3 cells

